# An integrated brain-machine interface platform with thousands of channels

**DOI:** 10.1101/703801

**Authors:** Elon Musk, Neuralink

## Abstract

Brain-machine interfaces (BMIs) hold promise for the restoration of sensory and motor function and the treatment of neurological disorders, but clinical BMIs have not yet been widely adopted, in part because modest channel counts have limited their potential. In this white paper, we describe Neuralink’s first steps toward a scalable high-bandwidth BMI system. We have built arrays of small and flexible electrode “threads”, with as many as 3,072 electrodes per array distributed across 96 threads. We have also built a neurosurgical robot capable of inserting six threads (192 electrodes) per minute. Each thread can be individually inserted into the brain with micron precision for avoidance of surface vasculature and targeting specific brain regions. The electrode array is packaged into a small implantable device that contains custom chips for low-power on-board amplification and digitization: the package for 3,072 channels occupies less than (23 × 18.5 × 2) mm^3^. A single USB-C cable provides full-bandwidth data streaming from the device, recording from all channels simultaneously. This system has achieved a spiking yield of up to 70% in chronically implanted electrodes. Neuralink’s approach to BMI has unprecedented packaging density and scalability in a clinically relevant package.

## 1 Introduction

Brain-machine interfaces (BMIs) have the potential to help people with a wide range of clinical disorders. For example, researchers have demonstrated human neuroprosthetic control of computer cursors [1, 2, 3], robotic limbs [4, 5], and speech synthesizers [6] using no more than 256 electrodes. While these successes suggest that high fidelity information transfer between brains and machines is possible, development of BMI has been critically limited by the inability to record from large numbers of neurons. Noninvasive approaches can record the average of millions of neurons through the skull, but this signal is distorted and nonspecific [7, 8]. Invasive electrodes placed on the surface of the cortex can record useful signals, but they are limited in that they average the activity of thousands of neurons and cannot record signals deep in the brain [9]. Most BMI’s have used invasive techniques because the most precise readout of neural representations requires recording single action potentials from neurons in distributed, functionally-linked ensembles [10].

Microelectrodes are the gold-standard technology for recording action potentials, but there has not been a clinically-translatable microelectrode technology for large-scale recordings [11]. This would require a system with material properties that provide high biocompatibility, safety, and longevity. Moreover, this device would also need a practical surgical approach and high-density, low-power electronics to ultimately facilitate fully-implanted wireless operation.

Most devices for long-term neural recording are arrays of electrodes made from rigid metals or semiconductors [12, 13, 14, 15, 16, 17, 18]. While rigid metal arrays facilitate penetrating the brain, the size, Young’s modulus and bending stiffness mismatches between stiff probes and brain tissue can drive immune responses that limit the function and longevity of these devices [19, 11]. Furthermore, the fixed geometry of these arrays constrains the populations of neurons that can be accessed, especially due to the presence of vasculature.

An alternative approach is to use thin, flexible multi-electrode polymer probes [20, 21]. The smaller size and increased flexibility of these probes should offer greater biocompatibility. However, a drawback of this approach is that thin polymer probes are not stiff enough to directly insert into the brain; their insertion must be facilitated by stiffeners [22, 21], injection [23, 24] or other approaches [25], all of which are quite slow [26, 27]. To satisfy the functional requirements for a high-bandwidth BMI, while taking advantage of the properties of thin-film devices, we developed a robotic approach, where large numbers of fine and flexible polymer probes are efficiently and independently inserted across multiple brain regions [28].

Here, we report Neuralink’s progress towards a flexible, scalable BMI that increases channel count by an order of magnitude over prior work. Our system has three main components: ultra-fine polymer probes (section 2 of this report), a neurosurgical robot (section 3), and custom high-density electronics (section 4). We demonstrate the rapid implantation of 96 polymer threads, each thread with 32 electrodes for a total of 3,072 electrodes.

We developed miniaturized custom electronics that allow us to stream full broadband electrophysiology data simultaneously from all these electrodes (section 5). We packaged this system for long-term implantation and developed custom online spike detection software that can detect action potentials with low latency. Together, this system serves as a state-of-the-art research platform and a first prototype towards a fully implantable human BMI.

## 2 Threads

We have developed a custom process to fabricate minimally displacive neural probes that employ a variety of biocompatible thin film materials. The main substrate and dielectric used in these probes is polyimide, which encapsulates a gold thin film trace. Each thin film array is composed of a “thread” area that features electrode contacts and traces and a “sensor” area where the thin film interfaces with custom chips that enable signal amplification and acquisition. A wafer-level microfabrication process enables high-throughput manufacturing of these devices. Ten thin film devices are patterned on a wafer, each with 3,072 electrode contacts.

Each array has 48 or 96 threads, each of those containing 32 independent electrodes. Integrated chips are bonded to the contacts on the sensor area of the thin film using a flip-chip bonding process. One goal of this approach is to maintain a small thread cross-sectional area to minimize tissue displacement in the brain. To achieve this, while keeping channel count high, stepper lithography and other microfabrication techniques are used to form the metal film at sub-micron resolution.

We have designed and manufactured over 20 different thread and electrode types into our arrays; two example designs are shown in panels A and B of fig. 1. Probes are designed either with the reference electrodes on separate threads or on the same threads as the recording electrodes (referred to as “on-probe references”). We have fabricated threads ranging from 5 to 50 μm in width that incorporate recording sites of several geometries (fig. 1). Thread thickness is nominally 4 to 6 μm, which includes up to three layers of insulation and two layers of conductor. Typical thread length is approximately 20 mm. To manage these long, thin threads prior to insertion, parylene-c is deposited onto the threads to form a film on which the threads remain attached until the surgical robot pulls them off. Each thread ends in a (16 × 50) μm^2^ loop to accommodate needle threading.

**Figure 1:**
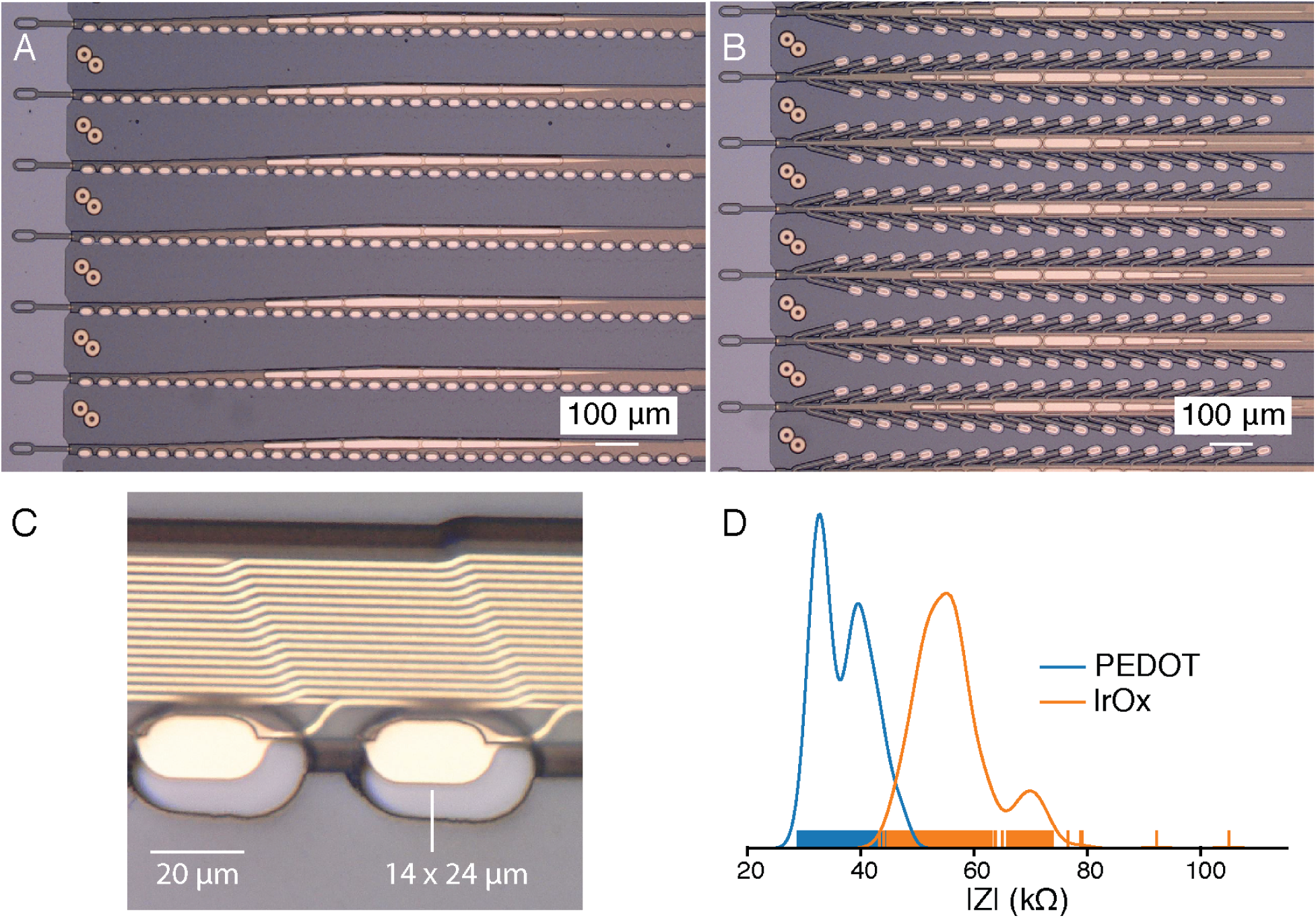
Our novel polymer probes. **A.** “Linear Edge” probes, with 32 electrode contacts spaced by 50 μm. **B.** “Tree” probes with 32 electrode contacts spaced by 75 μm. **C.** Increased magnification of individual electrodes for the thread design in panel A, emphasizing their small geometric surface area. **D.** Distribution of electrode impedances (measured at 1 kHz) for two surface treatments: PEDOT (*n* = 257) and IrOx (*n* = 588).

Since the individual gold electrode sites have small geometric surface areas (fig. 1C), we use surface modifications to lower the impedance for electrophysiology and increase the effective charge-carrying capacity of the interface (fig. 1D). Two such treatments that we have used are the electrically conductive polymer poly-ethylenedioxythiophene doped with polystyrene sulfonate (PEDOT:PSS) [29, 30] and iridium oxide (IrOx) [31, 32]. In bench-top testing we have achieved impedances of 36.97 ± 4.68 kΩ (*n* = 257 electrodes) and 56.46 ± 7.10 kΩ (*n* = 588) for PEDOT:PSS and IrOx, respectively. The lower impedance of PEDOT:PSS is promising, however the long-term stability and biocompatibility of PEDOT:PSS is less well established than for IrOx. These techniques and processes can be improved and further extended to other types of conductive electrode materials and coatings.

To keep the electronics package small, a novel alignment and flip-chip bonding process was developed. Multi-level gold stud bumps are placed throughout the PCB to act as alignment guides and temporary holders for the thin film. A custom shuttle is used to handle, align, and place the thin film on the PCB such that through holes in the thin film slide around the stud bumps. The thin film is secured into place by applying force to the gold stud bumps which flattens them into rivets. Next, the integrated chips are bonded directly both to contacts on the sensor area of the thin film and to pads on the PCB using standard flip-chip bonding processes. A custom silicon shuttle is used to vacuum-pick-up rows of 40 to 50 capacitors and bond a total of 192 capacitors onto the PCB. This alignment and bonding process was key to creating a package containing 3,072 channels in a (23 × 18.5) mm^2^ footprint.

## 3 Robot

Thin-film polymers have previously been used for electrode probes [21], but their low bending stiffness complicates insertions. Neuralink has developed a robotic insertion approach for inserting flexible probes [28], allowing rapid and reliable insertion of large numbers of polymer probes targeted to avoid vasculature and record from dispersed brain regions. The robot’s insertion head is mounted on 10 μm globally accurate, 400 mm × 400 mm × 150 mm travel three-axis stage, and holds a small, quick-swappable, “needle-pincher” assembly (fig. 2, fig. 3A).

**Figure 2:**
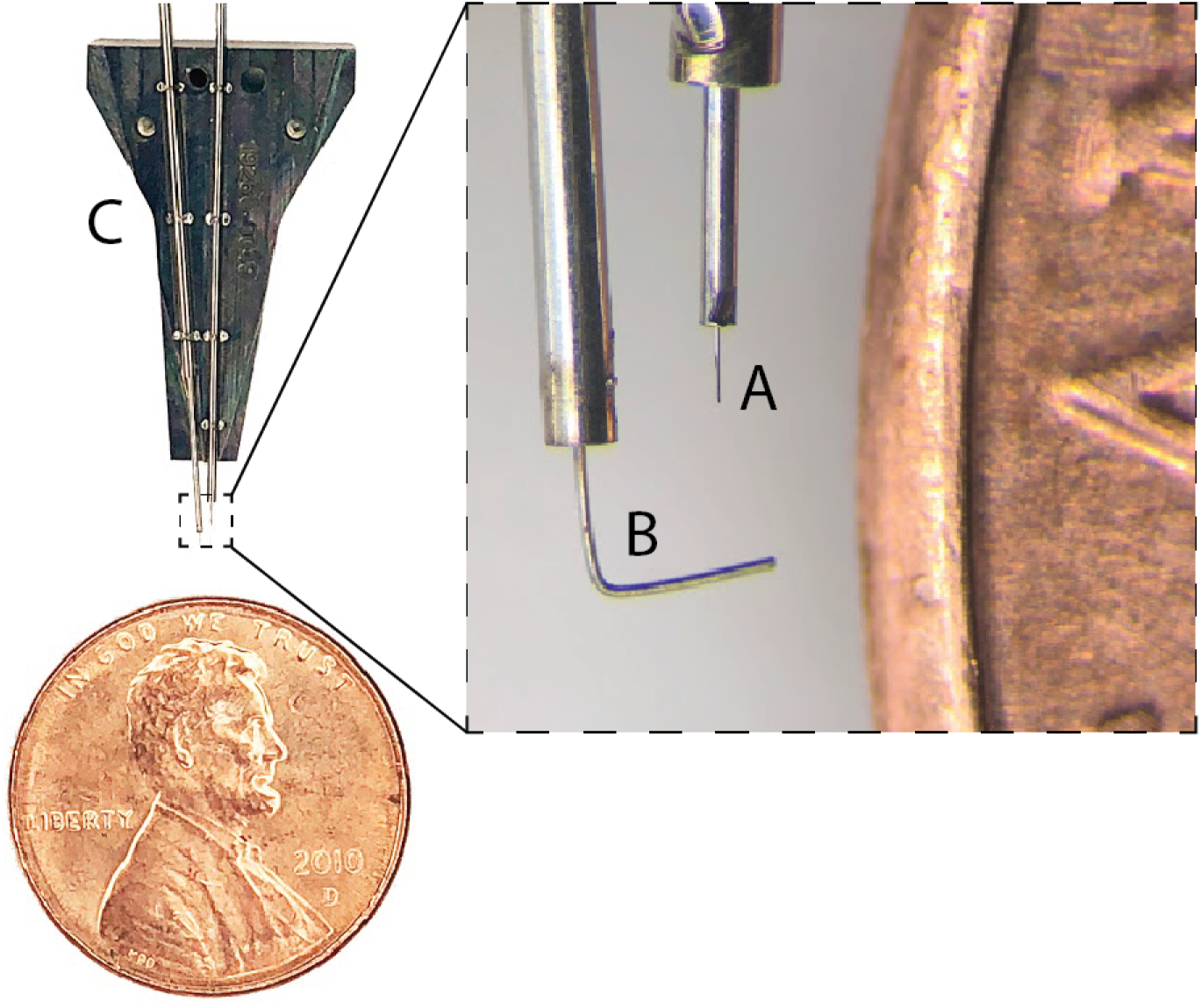
Needle pincher cartridge (NPC) compared with a penny for scale. **A.** Needle. **B.** Pincher. **C.** Cartridge.

**Figure 3:**
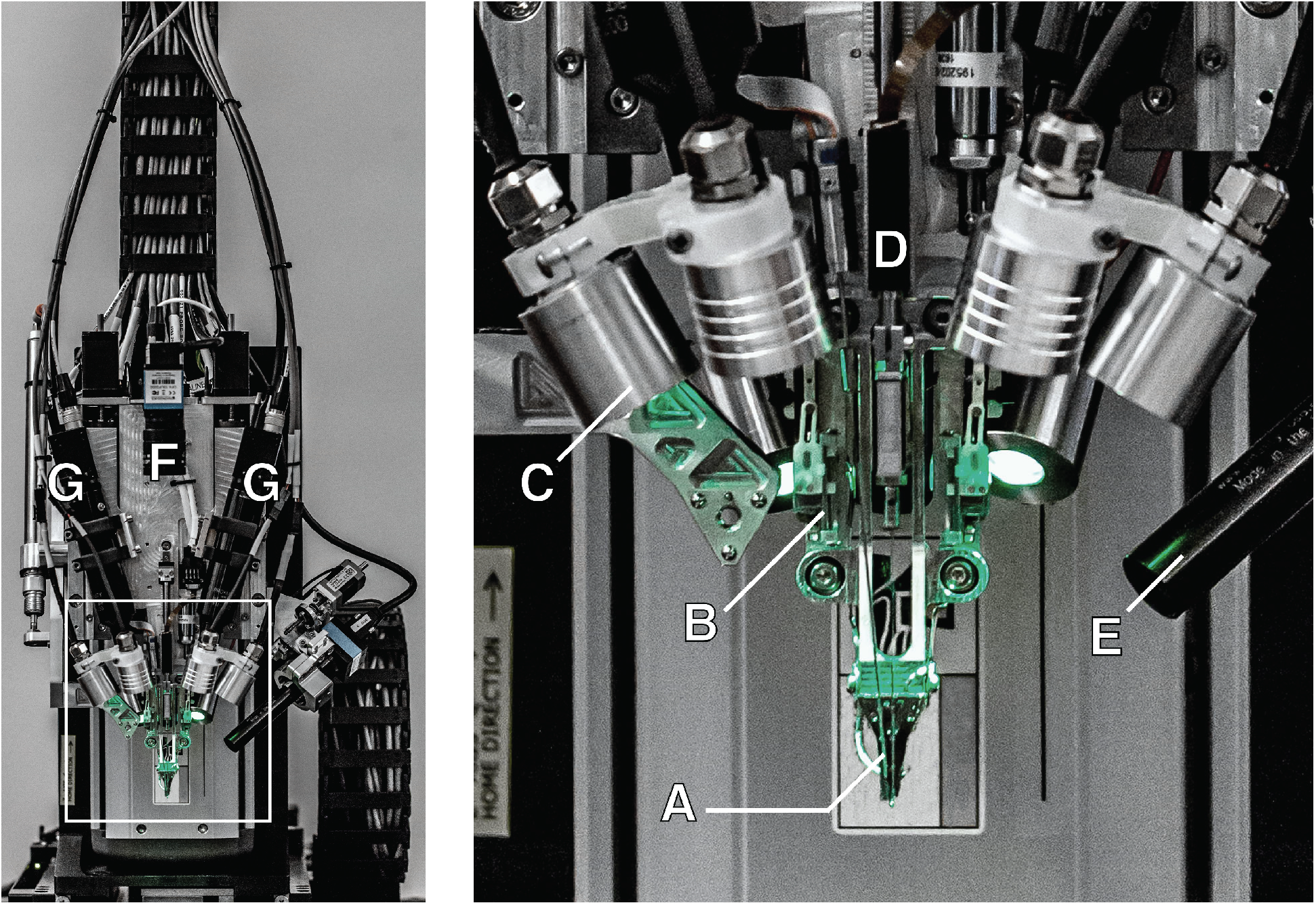
The robotic electrode inserter; enlarged view of the inserter-head shown in the inset. **A.** Loaded needle pincher cartridge. **B.** Low-force contact brain position sensor. **C.** Light modules with multiple independent wavelengths. **D.** Needle motor. **E.** One of four cameras focused on the needle during insertion. **F.** Camera with wide angle view of surgical field. **G.** Stereoscopic cameras.

The needle is milled from 40 μm diameter tungsten-rhenium wire-stock electrochemically etched to 24 μm diameter along the inserted length (fig. 2A). The tip of the needle is designed both to hook onto insertion loops—for transporting and inserting individual threads—and to penetrate the meninges and brain tissue. The needle is driven by a linear motor allowing variable insertion speeds and rapid retraction acceleration (up to 30,000 mm s^−2^) to encourage separation of the probe from the needle. The pincher is a 50 μm tungsten wire bent at the tip and driven both axially and rotationally (fig. 2B). It serves as support for probes during transport and as a guide to ensure that threads are inserted along the needle path. Figure 4 shows a sequence of photographs of the insertion process into an agarose brain proxy.

**Figure 4:**
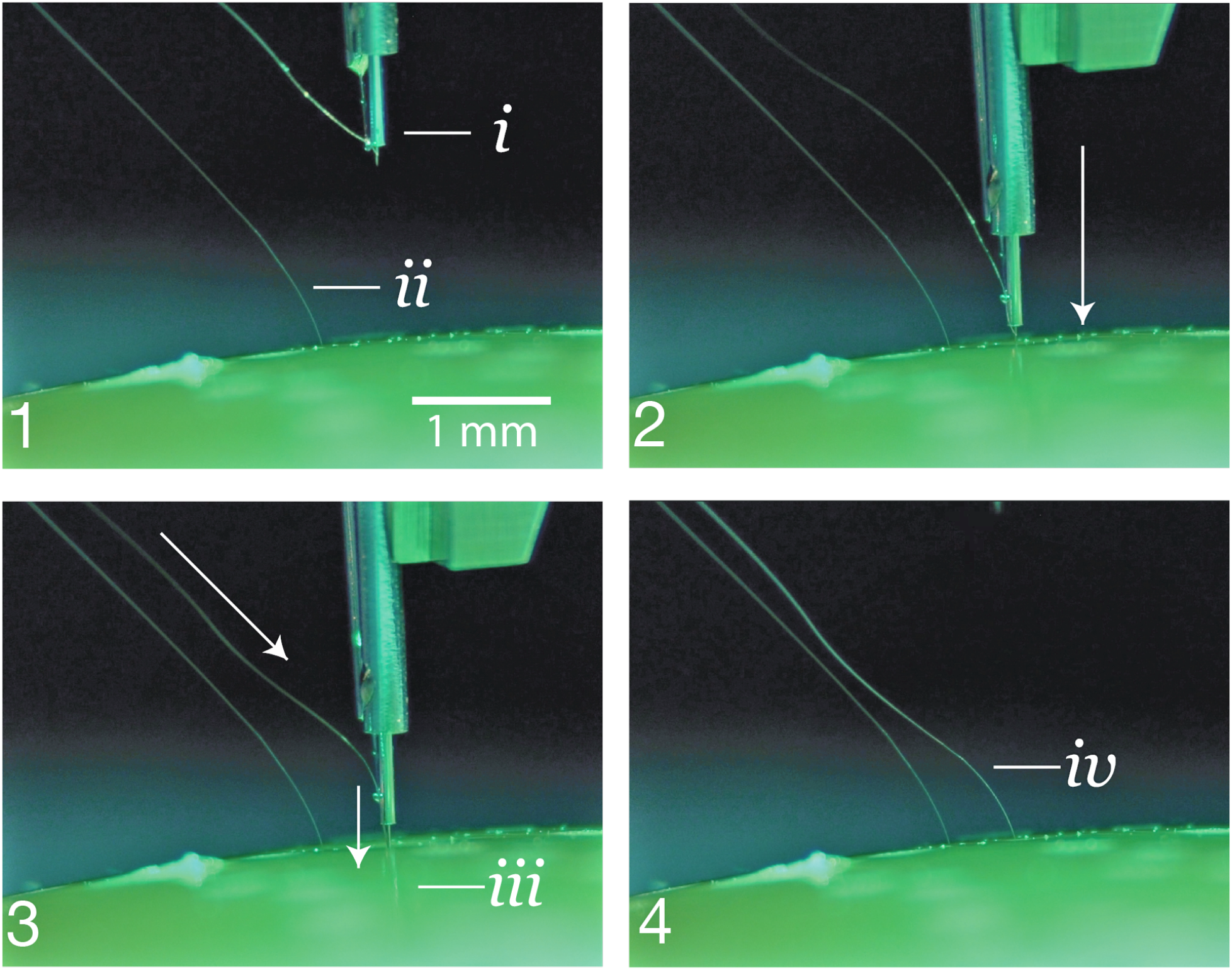
**1.** The inserter approaches the brain proxy with a thread. *i.* needle and cannula. *ii.* previously inserted thread. **2.** Inserter touches down on the brain proxy surface. **3**. Needle penetrates tissue proxy, advancing the thread to the desired depth. *iii.* inserting thread. **4.** Inserter pulls away, leaving the thread behind in the tissue proxy. *iv.* inserted thread.

The inserter head also holds an imaging stack (fig. 3E–G) used for guiding the needle into the thread loop, insertion targeting, live insertion viewing, and insertion verification. In addition, the inserter head contains six independent light modules, each capable of independently illuminating with 405 nm, 525 nm and 650 nm or white light (fig. 3C). The 405 nm illumination excites fluorescence from polyimide and allows the optical stack and computer vision to reliably localize the (16 × 50) μm^2^ thread loop and execute sub-micron visual servoing to guide, illuminated by 650 nm the needle through it. Stereoscopic cameras, software based monocular extended depth of field calculations, and illumination with 525 nm light allow for precise estimation of the location of the cortical surface.

The robot registers insertion sites to a common coordinate frame with landmarks on the skull, which, when combined with depth tracking, enables precise targeting of anatomically defined brain structures. An integrated custom software suite allows pre-selection of all insertion sites, enabling planning of insertion paths optimized to minimize tangling and strain on the threads. The planning feature highlights the ability to avoid vasculature during insertions, one of the key advantages of inserting electrodes individually. This is particularly important, since damage to the blood-brain barrier is thought to play a key role in the brain’s inflammatory response to foreign objects [33].

The robot features an auto-insertion mode, which can insert up to 6 threads (192 electrodes) per minute. While the entire insertion procedure can be automated, the surgeon retains full control, and if desired, can make manual micro-adjustments to the thread position before each insertion into the cortex. The neurosurgical robot is compatible with sterile shrouding, and has features to facilitate successful and rapid insertions such as automatic sterile ultrasonic cleaning of the needle. The needle pincher cartridge (NPC; fig. 2C) is the portion of the inserter head that makes direct contact with brain tissue and is a consumable that can be replaced mid-surgery in under a minute.

With this system, we have demonstrated an average of 87.1 ± 12.6 % (mean ± s.d.) insertion success rate over 19 surgeries. In this study, precise manual adjustments were made to avoid microvasculature on the cortical surface, slowing total insertion time from the fastest possible. Even with these adjustments, the total insertion time for this study averaged ~45 min, for an approximate insertion rate of 29.6 electrodes per minute (fig. 6). Insertions were made in a (4 × 7) mm^2^ bilateral craniotomy with >300 μm spacing between threads to maximize cortical coverage. This demonstrates that robotic insertion of thin polymer electrodes is an efficient and scalable approach for recording from large numbers of neurons in anatomically defined brain regions.

## 4 Electronics

Chronic recording from thousands of electrode sites presents significant electronics and packaging challenges. The density of recording channels necessitates placing the signal amplification and digitization stack within the array assembly, otherwise the cable and connector requirements would be prohibitive. This recording stack must amplify small neural signals (<10 μV_RMS_) while rejecting out-of-band noise, sample and digitize the amplified signals, and stream out the results for real-time processing—all using minimal power and size.

The electronics are built around our custom Neuralink application specific integrated circuit (ASIC), which consists of 256 individually programmable amplifiers (“analog pixels”), on-chip analog-to-digital converters (ADCs), and peripheral control circuitry for serializing the digitized outputs. The analog pixel is highly configurable: the gains and filter properties can be calibrated to account for variability in signal-quality due to process variations and the electrophysiological environment. The on-chip ADC samples at 19.3 kHz with 10 bit resolution. Each analog pixel consumes 5.2 μW and the whole ASIC consumes ~6 mW, including the clock drivers. Performance of the Neuralink ASIC is summarized in table 1 and a photograph of the fabricated device is shown in fig. 5A.

**Table 1:** Neuralink ASIC

|  |  |
| --- | --- |
| Number of Channels | 256 |
| Gain | 42.9–59.4 dB |
| Bandwidth | 3 Hz–27 kHz |
| Input-referred noise (3Hz - 10KHz) | 5.9 $\mu\text{V}_{\text{RMS}}$ |
| Maximum Differential Input Range | 7.2 mV<sub>PP</sub> |
| ADC resolution | 10 bit |
| Analog Pixel Power | 5.2 $\mu\text{W}$ |

**Figure 5:**
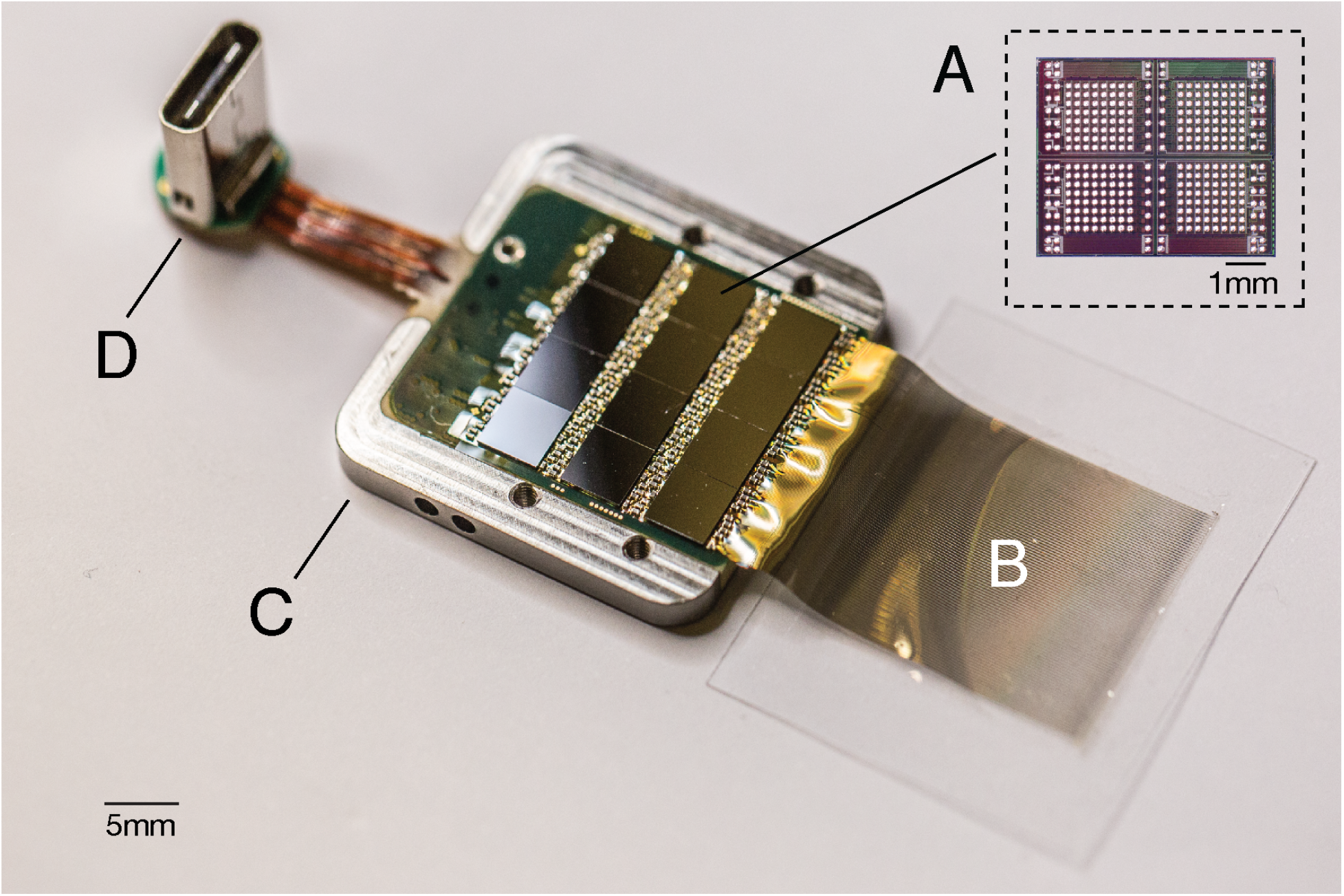
A packaged sensor device. **A.** individual neural processing ASIC capable of processing 256 channels of data. This particular packaged device contains 12 of these chips for a total of 3,072 channels. **B.** Polymer threads on parylene-c substrate. **C.** Titanium enclosure (lid removed). **D.** Digital USB-C connector for power and data.

**Figure 6:**
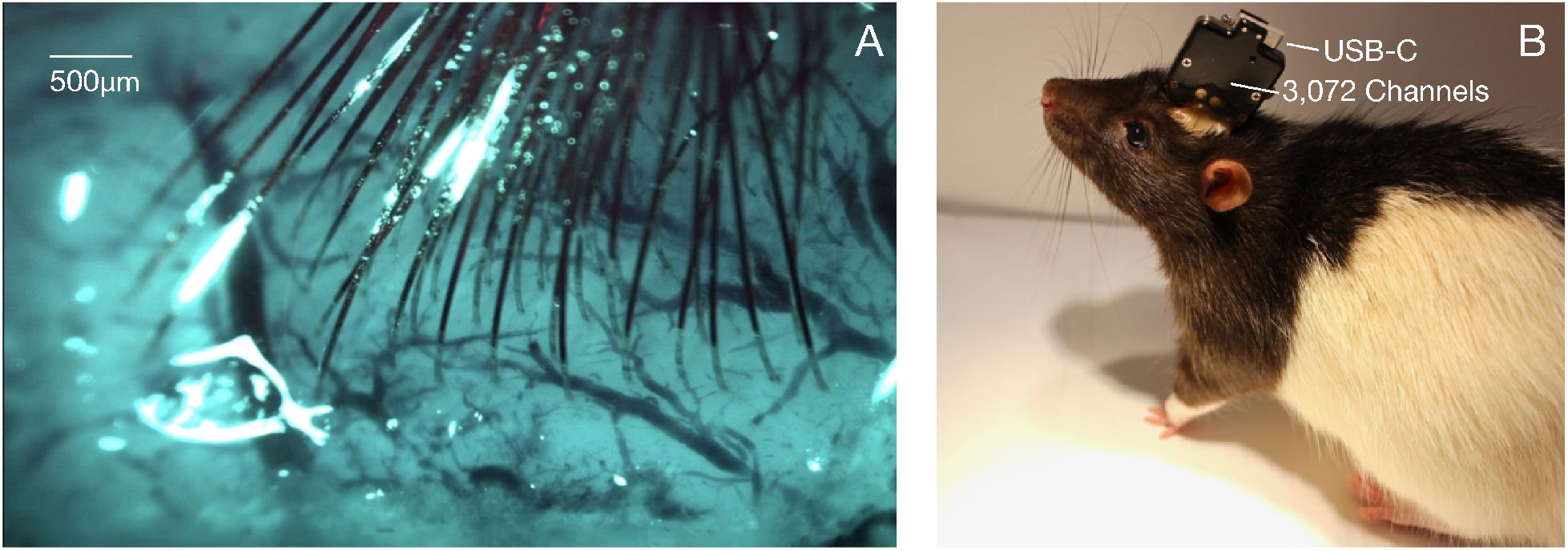
Thread implantation and packaging. **A.** An example peri-operative image showing the cortical surface with implanted threads and minimal bleeding. **B.** Packaged sensor device (“System B”) chronically implanted in a rat.

The Neuralink ASIC forms the core of a modular recording platform that allows for easy replacement of constitutive parts for research and development purposes (fig. 5). In the systems discussed here, a number of ASICs are integrated into a standard printed circuit board (PCB) using flip-chip integration. Each system consists of a field-programmable gate array (FPGA); real-time temperature, accelerometer, and magnetometer sensors; and a single USB-C connector for full-bandwidth data transfer. The systems are packaged in titanium cases which are coated with parylene-c, which serves as a moisture barrier to prevent fluid ingress and prolong functional lifetime.

We describe two such configurations that we have built, a 1,536 channel recording system (“System A”) and a 3,072 channel recording system (“System B”), summarized in table 2. While System A employs the current-generation Neuralink ASIC, System B uses an earlier revision with comparable functionality but poorer performance specifications. System B was designed to maximize channel density and is used for applications that demand extremely high channel count. In contrast, System A was designed to facilitate faster and more reliable manufacturing; it can be built five times faster than System B with better yields.

**Table 2:** Two recording system configurations

|  | System A | System B |
| --- | --- | --- |
| Number of Channels | 1,536 | 3,072 |
| Sampling Rate | 19.3 kHz | 18.6 kHz |
| Total System Power Consumption | 550 mW | 750 mW |
| Total System Size | $(24.5 \times 20 \times 1.65) \text{ mm}^3$ | $(23 \times 18.5 \times 2) \text{ mm}^3$ |
| Implant Weight | 11 g | 15 g |

An ethernet-connected base station converts the data streams from these systems into multicast 10G ethernet UDP packets allowing downstream users to process the data in a variety of ways, e.g. visualizing the data in real-time [34] or writing it to disk. Each base station can connect to up to three implants simultaneously. These devices are further supported by a software ecosystem that allows for plug and play usability with zero configuration: neural data begins streaming automatically when a cable is connected.

## 5 Electrophysiology

We have implanted both systems A and B in male Long-Evans rats, as described in section 3. All animal procedures were performed in accordance with the National Research Council’s *Guide for the Care and Use of Laboratory Animals* and were approved by the Neuralink Institutional Animal Care and Use Committee. Electrophysiological recordings were made as the animals freely explored an arena equipped with a commutated cable that permitted unrestricted movement. System A can record 1,344 out of 1,536 channels simultaneously, the exact channel configuration can be arbitrarily specified at the time of recording; system B can record from all 3,072 channels simultaneously. Digitized broadband signals were processed in real-time to identify action potentials (spikes) using an online detection algorithm.

Spike detection requirements for real-time BMI are different from most conventional neurophysiology. While most electrophysiologists spike-sort data offline and spend significant effort to reject false-positive spike events, BMI events must be detected in real time and spike detection parameters must maximize decoding efficacy. Using our custom online spike-detection software, we found that a permissive filter that allows an estimated false positive rate of ~0.2 Hz performs better than setting stringent thresholds that may reject real spikes (data not shown).

Given these considerations, we set a threshold of >0.35 Hz to quantify the number of electrodes that recorded spiking units. Since we typically do not spike sort our data, we do not report multiple units per channel. BMI decoders commonly operate without spike sorting with minimal loss of performance [36, 37]. Moreover, recent results show that spike sorting is not necessary to accurately estimate neural population dynamics [38].

Data from a recent experiment using System A are shown in fig. 7 and fig. 8. In this experiment, 40 of 44 attempted insertions were successful (90 %) for a total of 1280 implanted electrodes, of which 1,020 were recorded simultaneously. The broadband signals recorded from a representative thread show both local field and spiking activity fig. 7. A sample output of the spike detection pipeline is shown in raster form in fig. 8. In this example, two overlapping recording configurations were used to record from all 1,280 implanted channels. On this array, our spiking yield was 43.4 % of channels, with many spikes appearing on multiple neighboring channels, as has been observed in other high-density probes [16, 17, 21]. On other System A arrays we observed 45.60 ± 0.03 % (mean ± s.e.m.) across 19 surgeries with a maximum spiking yield of 70 %.

**Figure 7:**
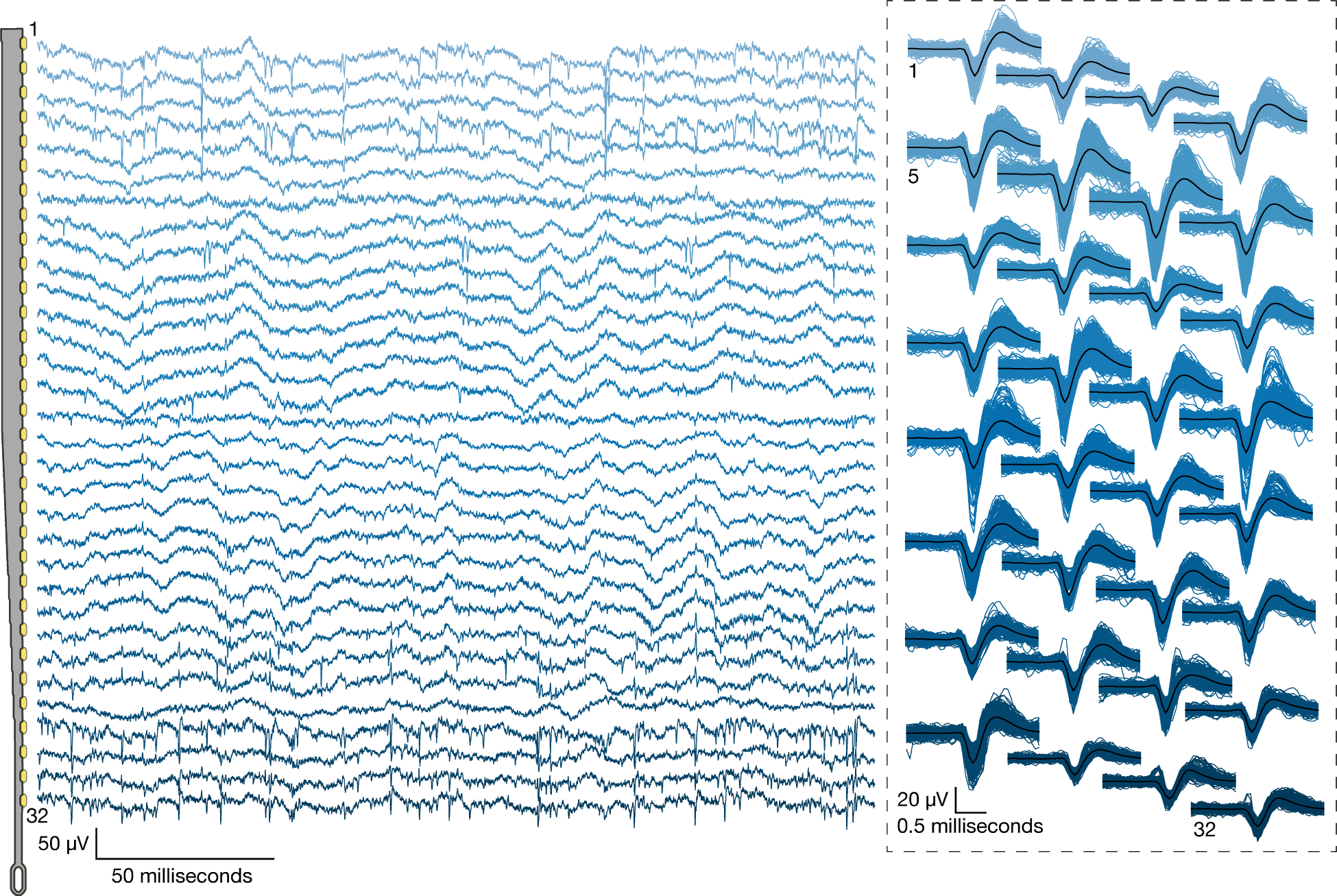
**Left:** Broadband neural signals (unfiltered) simultaneously acquired from a single thread (32 channels) implanted in rat cerebral cortex. Each channel (row) corresponds to an electrode site on the thread (schematic at left; sites spaced by 50 μm). Spikes and local field potentials are readily apparent. **Right:** Putative waveforms (unsorted); numbers indicate channel location on thread. Mean waveform is shown in black.

**Figure 8:**
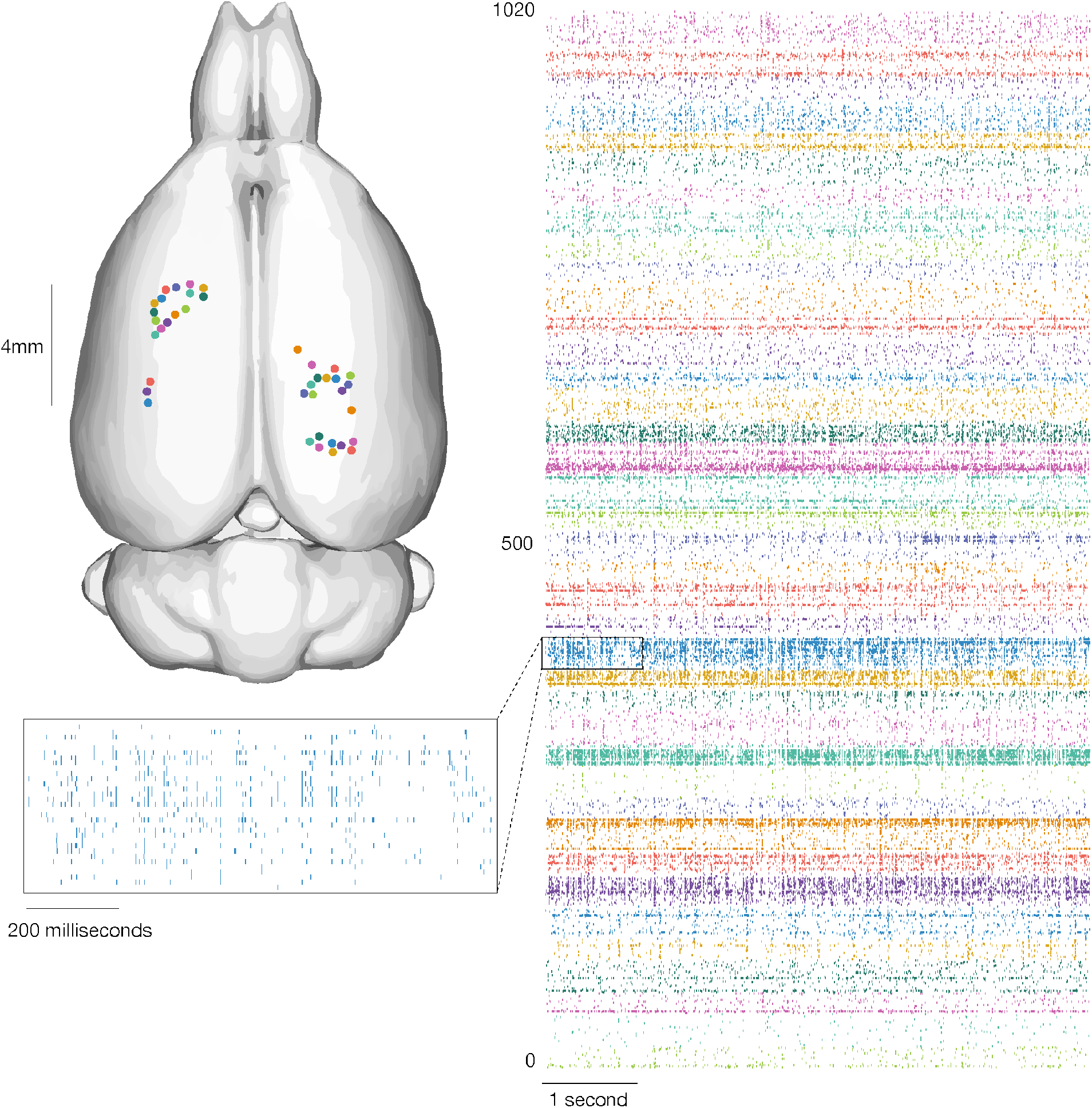
Our devices allow the recording of widespread neural activity distributed across multiple brain regions and cortical layers. **Left:** thread insertion sites (colored circles) are indicated on rendered rodent brain. [35] **Right:** raster of 1,020 simultaneously recorded channels, sorted per thread (color corresponds to insertion site). **Inset:** enlarged raster of spikes from a single thread. This thread corresponds to the one shown in fig. 7.

## 6 Discussion

We have described a BMI with high-channel count and single-spike resolution. It is based on flexible polymer probes, a robotic insertion system, and custom low-power electronics. This system serves two main purposes: it is a research platform for use in rodents and serves as a prototype for future human clinical implants. The ability to quickly iterate designs and testing in rodents allows for the rapid refinement of devices, manufacturing processes, and software. Because it is a research platform, the system uses a wired connection to maximize the bandwidth for raw data streaming. This is important for performance assessments and crucial for the development of signal processing and decoding algorithms. In contrast, the clinical devices that will derive from this platform will be fully implantable—which requires hermetic packaging—and have on-board signal compression, reduced power consumption, wireless power transmission, and data telemetry through the skin without percutaneous leads.

Modulating neural activity will be an important part of next-generation clinical brain-machine interfaces [39], for example to provide a sense of touch or proprioception to neuroprosthetic movement control [40, 41]. Therefore, we designed the Neuralink ASIC to be capable of electrical stimulation on every channel, although we have not demonstrated these capabilities here.

This BMI system has several advantages over previous approaches. The size and composition of the thin-film probes are a better match for the material properties of brain tissue than commonly used silicon probes, and therefore may exhibit enhanced biocompatibility [28, 21]. Also, the ability to choose where our probes are inserted, including into sub-cortical structures, allows us to create custom array geometries for targeting specific brain regions while avoiding vasculature. This feature is significant for creating a high-performance BMI, as the distribution of electrodes can be customized depending on the task requirements. Lastly, the miniturization and design of the Neuralink ASIC affords great flexibility in system design and supports very high channel counts within practical size and power constraints.

In principle, our approach to brain-machine interfaces is highly extensible and scalable. Here we report simultaneous broadband recording from over 3,000 inserted electrodes in a freely moving rat. In a larger brain, multiple devices with this architecture could be readily implanted, and we could therefore interface with many more neurons without extensive re-engineering. Further development of surgical robotics could allow us to accomplish this without dramatically increasing surgery time.

While significant technological challenges must be addressed before a high-bandwidth device is suitable for clinical application, with such a device, it is plausible to imagine that a patient with spinal cord injury could dexterously control a digital mouse and keyboard. When combined with rapidly improving spinal stimulation techniques [42], in the future this approach could conceivably restore motor function. High-bandwidth neural interfaces should enable a variety of novel therapeutic possibilities.

## 7 Acknowledgments

We would like to acknowledge the contributions of Lawrence Livermore National Laboratory (LLNL), Berkeley Marvell Nanofabrication Laboratory, Berkeley Wireless Research Center (BWRC), Stanford Nanofabrication Facility, and former and current Neuralink employees to the work described here. Thread manufacturing and high density package assembly was performed under a collaboration with LLNL (CRADA No. TC02267).

## 8 Supplementary Videos

**Video 1:** A series of six insertions by the neurosurgical robot into an agarose brain proxy. Thread-capture by the needle occurs off-frame. The changes background color are caused by illumination with different frequencies of light at different stages of the threading and insertion process. One thread was inserted before the start of the video.

**Video 2:** A 3D rendered view of thread arrangement (same data presented in fig. 8). Thread insertion is visualized in the same order as in the actual surgery, but time has been compressed for presentation. Thread size and insertion depth are representative. The stereotaxic coordinates of each insertion are represented on the dataset provided by Calabrese and coworkers [35].

